# *Ab initio* CryoEM maps minimizing preferred orientation problems in heterogeneous mixtures: ReconSIREN

**DOI:** 10.64898/2026.10.03.756391

**Authors:** D. Herreros, C.O.S. Sorzano, J.M. Carazo

**Affiliations:** Centro Nacional de Biotecnología-CSIC, C/ Darwin, 3, 28049, Cantoblanco, Madrid, Spain

**Keywords:** Cryo-Electron Microscopy (Cryo-EM), Deep Learning, *ab initio* reconstruction, amortized inference

## Abstract

Cryo-Electron Microscopy (CryoEM) Single Particle Analysis (SPA) is pivotal for elucidating protein structures and their conformational dynamics. Several critical steps are required to obtain accurate maps. First, the *ab initio* relative orientations and shifts between images must be determined. Second, the common issue of specimens presenting a preferred orientation should be adequately addressed. Third, all of the above must account for particle flexibility and sample heterogeneity. Additionally, these steps are interrelated, making the search for all these parameters a real challenge when considering convergence robustness. In this context, we present ReconSIREN, a novel deep learning framework that addresses the key steps outlined before in an *ab initio* manner while greatly minimizing one of the most important practical burdens in SPA, namely the errors introduced in the CryoEM maps by the existence of preferred orientations; the latter is achieved by a change of the optimization approach that avoids the so-called “attraction problem”, and by the introduction of a new type of geometrical priors. It does so fully *ab initio*, using deep learning training/inference steps without resorting to more traditional optimization approaches, and generating highly accurate initial reconstructions. We present maps as both homogeneous (consensus) maps and a full conformational landscape for analyzing flexibility and heterogeneity. Thanks to its Mixture-of-Experts (MoE) architecture, ReconSIREN distributes the estimation workload across independent experts, improving optimization efficiency and simplifying the process.

## Introduction

CryoEM Single Particle Analysis (SPA) has become a leading technique for studying protein structure and conformational heterogeneity, thanks to its ability to capture individual images of biological macromolecules [1]. This enables image-processing workflows to perform individualized image analysis and recover key parameters that enable the retrieval of conformational ensembles [2, 3].

In any CryoEM image-processing workflow, determining the initial volume for final refinement, along with the associated angular assignments and in-plane shifts, is critical, as these estimates form the basis for subsequent processing steps [4]. An error at this stage will most likely lead to a local minimum that may represent an artefactual map. Consequently, the *ab initio* estimation of these parameters should be sufficiently accurate to aid in finding an accurate final reconstruction. Additionally, these parameters are critical for any heterogeneity analysis algorithm, as most approaches assume that the poses are either fixed or very close to a good local minimum [5].

A key consideration is that we designed ReconSIREN to tackle one of the fundamental problems in CryoEM: estimating correct maps in the presence of strong preferred orientations, a situation that affects a large percentage of practical cases and remains a classic bottleneck for accurate structural resolution of many specimens [6]. At this point, it should be remembered that the presence of preferred orientations in a data set is not necessarily a source of map artifacts in itself [7]. Preferred orientation often does not mean unique orientation, but rather a different proportion of views in different projection directions. This imbalance usually becomes an artifact because of the so-called “attraction problem”, first identified in [8] and since then acknowledged many times [9–11]. In short, geometry imbalance generally induces images to be drawn towards projection directions with a higher Signal-to-Noise Ratio (SNR), so that the angular assignment of minoritarian classes is wrong. This incorrect assignment is the main cause of artifacts in the reconstructions, significantly limiting the ability to recover high-resolution structural features and interpret the resulting volumes. In this work, we will present striking examples of this problem and how ReconSIREN greatly minimizes them.

These examples also show that accuracy is not the only requirement for *ab initio* parameter estimation; robustness is essential. Ideally, parameter estimation should always converge to the correct solution with the highest possible quality. Given the complex conditions of CryoEM images (low SNR, preferred orientations, heterogeneity…), convergence across different experiments remains a challenge that is not fully resolved. ReconSIREN’s design also improves the method’s convergence, aiming to provide robust *ab initio* estimates across proteins with varying characteristics.

ReconSIREN introduces a novel pose-ensemble optimization that leverages *a priori* information about the ensemble angular distribution to promote correct image-to-projection assignments while mitigating the tendency to be drawn toward high-SNR projection directions. As a result, ReconSIREN reports more accurate angular assignments and *ab initio* reconstructions while improving convergence and without attraction-derived artifacts.

While ReconSIREN presents a novel approach to the complex case of *ab initio* estimation of angular/shift assignments, reconstruction, and heterogeneity, several previous works have partially addressed these issues, as discussed below.

To further improve the quality of *ab initio* reconstructions, some approaches have tackled the problem using deep learning (*e.g.*, CryoAI [12], CryoSPIN [13]; for a review, see [14] and the references therein). These approaches employed a neural network to jointly estimate the projection direction of the images and the *ab initio* homogeneous reconstruction. However, because they rely on local searches to explore a complex solution space, these methods require some *a priori* information about the images (*e.g.*, properly centering them beforehand), which prevents them from being applied in a fully *ab initio* manner.

Following the previous principles, methods such as CryoFIRE [15] adapted neural networks to include *ab initio* estimation of conformational landscapes and heterogeneous states. However, these approaches still suffered from limitations similar to those of the previously introduced methods and required *a priori* information about the images.

To simplify image-parameter optimization, recent approaches in this area, such as CryoDRGN-AI [16], combine a classical exhaustive grid search with local optimization, enabling them to estimate angular assignments, in-plane shifts, and heterogeneous states *ab initio* from the images. However, adding a classical step to the optimization process prevents full amortization of the estimated parameters, a key advantage of deep learning over classical methods.

In this rich context of previous work, ReconSIREN achieves its uniqueness through several novelties, building on the many innovations already present in HetSIREN [17]. Architecturally, ReconSIREN uses a Mixture-of-Experts (MoE) model. Our MoE lets the network train independent experts that estimate only a subset of image parameters, making optimization more efficient and simpler. Then, at the level of priors, ReconSIREN introduces a novel pose-ensemble optimization that leverages *a priori* information about the ensemble angular distribution to promote the correct assignment of images to a projection direction while mitigating the tendency to be drawn toward high-SNR projection directions. As a result, ReconSIREN can converge towards more accurate angular assignments and high-resolution *ab initio* reconstructions without attraction-derived artifacts in a full deep learning approach, avoiding, in most cases, trapping the final map in an artefactual local minimum. Naturally, after initial model building and heterogeneity detection, ReconSIREN can be complemented with other methods, including repicking original micrographs to increase the proportion of minoritarian views, other approaches to refine projection directions, or procedures to extrapolate views into missing angular regions.

## Results

To simplify the discussion of the results obtained with ReconSIREN, their presentation will be divided into four different cases:

1. Evaluation of the alignments and consensus volume accuracy from experimental images.
2. Mitigation of attraction due to the presence of preferred orientations.
3. Evaluation of the conformational spaces derived from experimental images.
4. Evaluation of ReconSIREN when 2D averages are used to train the Mixture of Experts (MoE) model.
5. Analysis of the maximum resolution achievable by ReconSIREN.

In all cases, ReconSIREN was configured to estimate alignments, in-plane shifts, consensus volume, and the conformational landscape fully *ab initio*. The methods section describes alternative ways to execute ReconSIREN, depending on the prior information available in the data.

ReconSIREN is freely distributed as part of the package *hax-em* as described in the Code Availability section, simplifying its installation and application.

### Evaluation of ReconSIREN alignments and consensus volumes derived from experimental data

One aim of ReconSIREN is to find the *ab initio* angular assignments and in-plane shifts from a particle dataset, as well as the consensus reconstruction derived from the images. Because this is a strongly coupled problem, ReconSIREN uses two MoE models: the first finds the images’ angular assignments and in-plane shifts, while the second determines the consensus map. In this way, each model works independently on its own problem but can leverage the other model’s improvements to find a better solution.

To validate the accuracy and performance of ReconSIREN in solving the previous two problems, we evaluated the method on five different datasets with varying properties:

- PKM2 [18]: This dataset exhibits strong rigidity and a mid-preferred orientation issue. The dataset comprises ∼120k images with a box size of 256 pixels and a sampling rate of 1.00 Å/px.
- SARS-Cov-2 spike: This dataset captures the down-to-up motion of the spike RBD. The dataset comprises ∼200k images with a box size of 224 pixels and a sampling rate of 1.25 Å/px.
- GR:Hsp90:FKBP51 complex [19]: This dataset shows strong flexibility in the GR region of the complex. The dataset comprises ∼120k images with a box size of 320 pixels and a sampling rate of 0.83 Å/px.
- HER2 in complex with Trastuzumab (in amphipols) [20]: This dataset is challenging, especially because of the very complex flexibility pattern of the complete protein, which has only one transmembrane helix. Accordingly, we focused only on the extracellular domain, which has its own flexibility pattern and a total mass of around 80 kDa. Additionally, the dataset presents preferred orientations, making it harder to process. The next section explains how ReconSIREN addresses preferred orientations in the dataset, which was also important to achieve the HER2 results presented here. It comprises ∼100k images with a box size of 200 pixels and a sampling rate of 1.30 Å/px.
- ABCB11: This dataset is challenging not only due to the flexibility of the protein but also due to the presence of a nanodisc, which also induces preferred orientations. It comprises ∼100k images with a box size of 200 pixels and a sampling rate of 1.09 Å/px.

We chose these datasets because they provide a good sample for evaluating the method’s performance on proteins with varying characteristics and experimental conditions.

As an initial experiment, we trained an independent ReconSIREN network on the raw particle images (downsampled to a 128-pixel box size internally to improve training performance), with only the Contrast Transfer Function (CTF) pre-estimated. Downsampling the images to a 128-pixel box size is one way to run ReconSIREN, where resolution is sacrificed for higher performance. We recommend this setup when lower training times are desirable, as it yields fast *ab initio* estimates to use later in an image processing workflow. However, this is not the only way to run ReconSIREN; later we show how to run the software to maximize resolution and the accuracy of the estimated parameters and maps.

Figure 1 shows the learned angular assignments and consensus volumes. ReconSIREN also had to predict the in-plane shifts from scratch, without assuming any pre-centering, which made the optimization more challenging. The angular distributions presented in this work are shown on a logarithmic scale, which simplifies visual assessment of the image distribution in the projection sphere.

**Figure 1:**
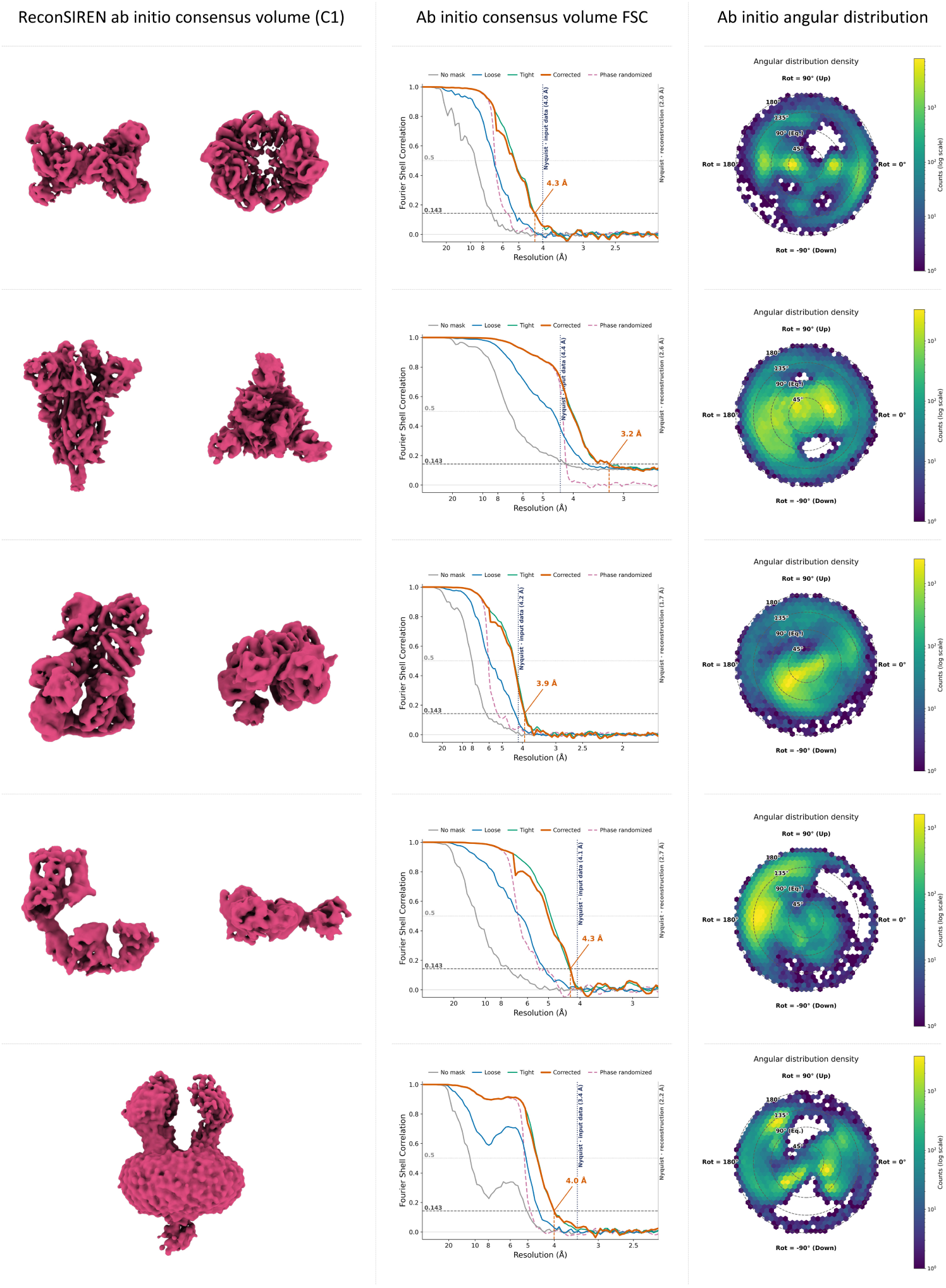
Example of the *ab initio* consensus reconstructions, FSC curves, and angular assignments obtained with ReconSIREN for five different datasets (from top to bottom - PKM2, SARS-CoV-2 spike, GR:Hsp90:FKBP51 complex, HER2, ABCB11) using experimental particle images. For the ABCB11 case, we present only the side view of the map, as the top view was not very informative. We did not consider symmetry in any of the datasets presented. For each macromolecule, we show two views of the reconstructed map, the FSC, and its angular distribution.

In Supplementary Figure 1, we compare logarithmic and linear scales for the distributions shown in Figure 1 to better assess the scale’s effect on visualization.

As shown in the Figure, ReconSIREN can learn high-quality *ab initio* maps and angular assignments without apparent attraction or preferred-orientation issues. In the next section, we will further discuss the method’s robustness to preferred orientation and its significant mitigation of attraction artifacts. Although the network can handle symmetry, we obtained all examples without assuming any symmetry.

Even when resolution is limited by image size, ReconSIREN can recover *ab initio* volumes and image alignments that retain high-resolution features compared with standard initial volume programs. Therefore, ReconSIREN estimations provide a better starting point for subsequent steps in an image processing workflow, which may aid in the search for more accurate parameters and structures.

To better evaluate the quality of the *ab initio* consensus maps, we used the network-derived angular assignments and in-plane shifts as input to a CryoSPARC [21] homogeneous reconstruction job to estimate their FSC. For this step, we reassigned the estimated angular assignments and in-plane shifts to the full-resolution images to show that standard software can further work and benefit from ReconSIREN estimates. Therefore, the homogeneous reconstructed maps have a lower Nyquist resolution than the network observes. To make the FSC plots included in Figure 1 more informative, we have also included the Nyquist resolution observed by the network.

### Avoiding preferred orientation attraction on angular assignments with ReconSIREN

In addition to building ReconSIREN to accurately estimate the consensus map and image alignments, we paid close attention to the attraction problem that CryoEM suffers when aligning images to build a density map.

Attraction artifacts arise because, when aligning images with a low signal-to-noise ratio (SNR), projection directions that produce stronger signals tend to attract those images in which the signal features are less prominent, even when the true projection directions of those images differ substantially [8–10].

Consequently, preferred orientations in cryo-EM datasets exacerbate this effect by creating projection directions with systematically higher SNR, making them disproportionately attractive during alignment.

Preferred orientations and attraction artifacts remain among the main limitations of CryoEM for achieving high resolution in many important biological specimens. Although methods such as spIsonet [22] have addressed this problem by training a network to correct maps affected by these artifacts, the underlying mathematical problem of proper angular assignment remains largely unresolved. Therefore, in ReconSIREN, we address the fundamental issue of properly assigning projection directions while avoiding attraction by using network ensembles that predict uniform particle poses, which largely minimizes the chance of falling into a local minimum during optimization. Thanks to these ensembles, the network can see, at every training step, a wide set of plausible projection directions, allowing it to determine which is most appropriate for a given image based on the consensus map. The consensus map also benefits from the ensemble-based pose assignment, allowing it to converge towards a more isotropic volume. The Methods section describes how these ensembles work.

To evaluate the robustness of ReconSIREN against datasets exhibiting strong preferred orientations, we analyze two representative datasets:

- Pose-entangled dataset from the 2026 CAHRA challenge [23]: This simulated dataset contains images of a channel protein with C11 symmetry. The dataset was deliberately simulated to have a strong preferred orientation and two conformational states with different populations. The dataset comprises ∼30k images with a box size of 256 pixels, a sampling rate of 1.20 Å/px, CTF, and realistic noise.
- Hemagglutinin from EMPIAR 10096 (untilted acquisition) [10]. This dataset is one of the best-known examples of preferred orientation in EMPIAR. For our experiments, we downloaded the deposited, untilted particle images and kept only the CTF information. We did not follow any repicking strategy to increase the number of side views in the dataset; instead, we used only the (untilted) deposited images to better evaluate ReconSIREN’s performance on a challenging dataset.

First, we evaluated the quality of the consensus *ab initio* reconstructions as in the previous sections. The results from these experiments are summarized in Figure 2. As in the previous section, we assumed no symmetry during ReconSIREN training, so all results presented correspond to C1 point-group symmetry.

**Figure 2:**
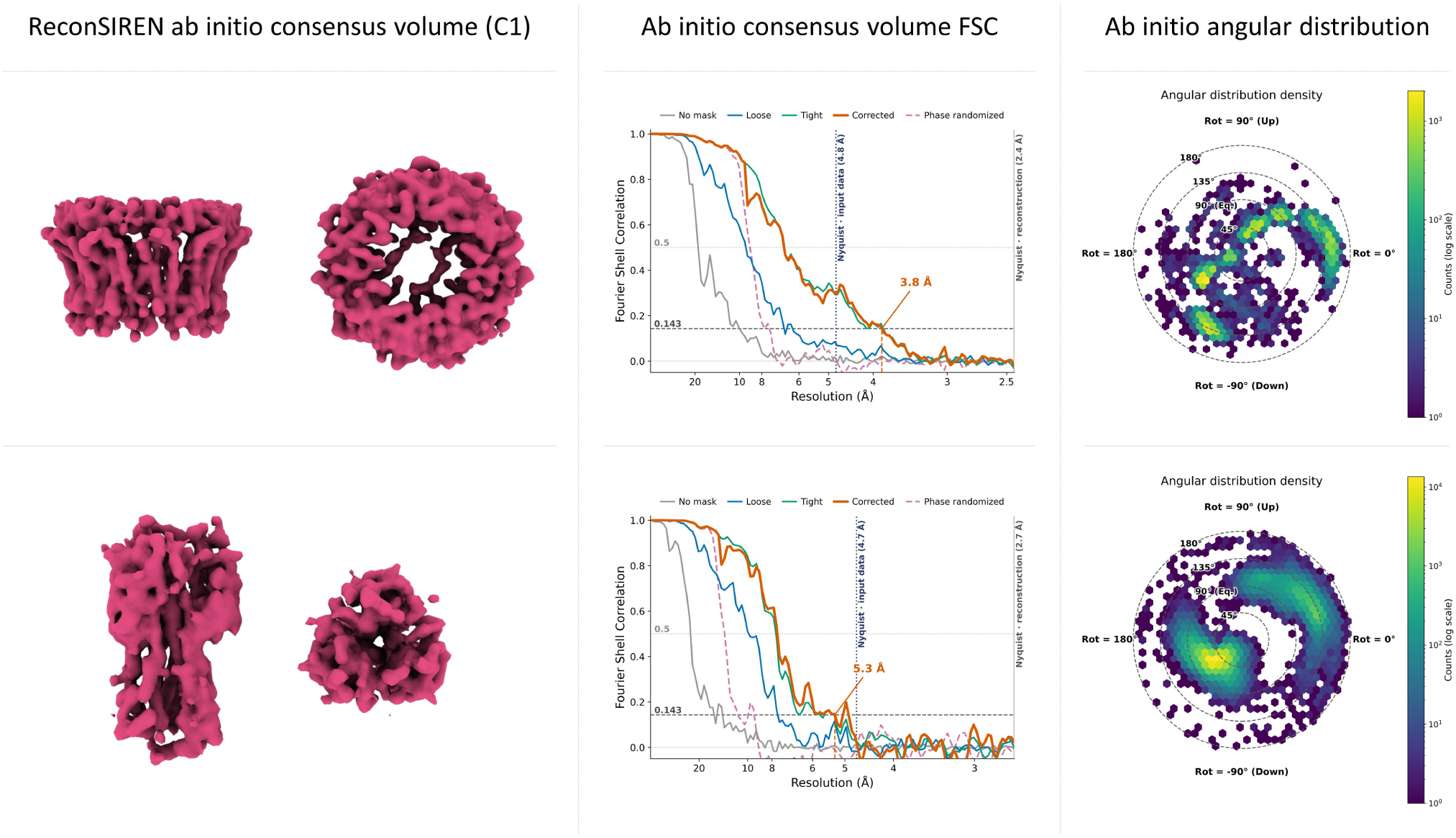
Example of the *ab initio* consensus reconstructions, FSC curves, and angular assignments obtained with ReconSIREN for two datasets with a strong preferential orientation (from top to bottom: CAHRA pose entanglement dataset, hemagglutinin dataset from EMPIAR 10096. We did not consider symmetry in any of the datasets presented. As the results show, ReconSIREN learns an angular distribution that strongly mitigates attraction induced by the data’s preferred orientation, thereby significantly improving the quality of the *ab initio* reconstruction.

As shown in the network, ReconSIREN reconstructs accurate *ab initio* maps whose FSCs follow a trend similar to that presented in the previous section. Similar to the previous experiments, we downsampled the training images to a box size of 128 pixels, whereas we computed the FSCs with CryoSPARC without downsampling. This allows us to show how to couple ReconSIREN results with 3D refinement programs, as shown later in this section.

A more interesting result emerges when comparing ReconSIREN’s performance with other widely used *ab initio* reconstruction methods, such as CryoSPARC. Figure 3a shows the *ab initio* volumes obtained from these two methods when no symmetry is assumed. The comparison clearly shows that, because of the data’s strong preferred orientation and the protein’s high symmetry, CryoSPARC fails to retrieve a good *ab initio* volume, which is strongly affected by attraction artifacts. In contrast, ReconSIREN’s robustness to attraction enables the network to produce a high-resolution *ab initio* map even without symmetry considerations.

**Figure 3:**
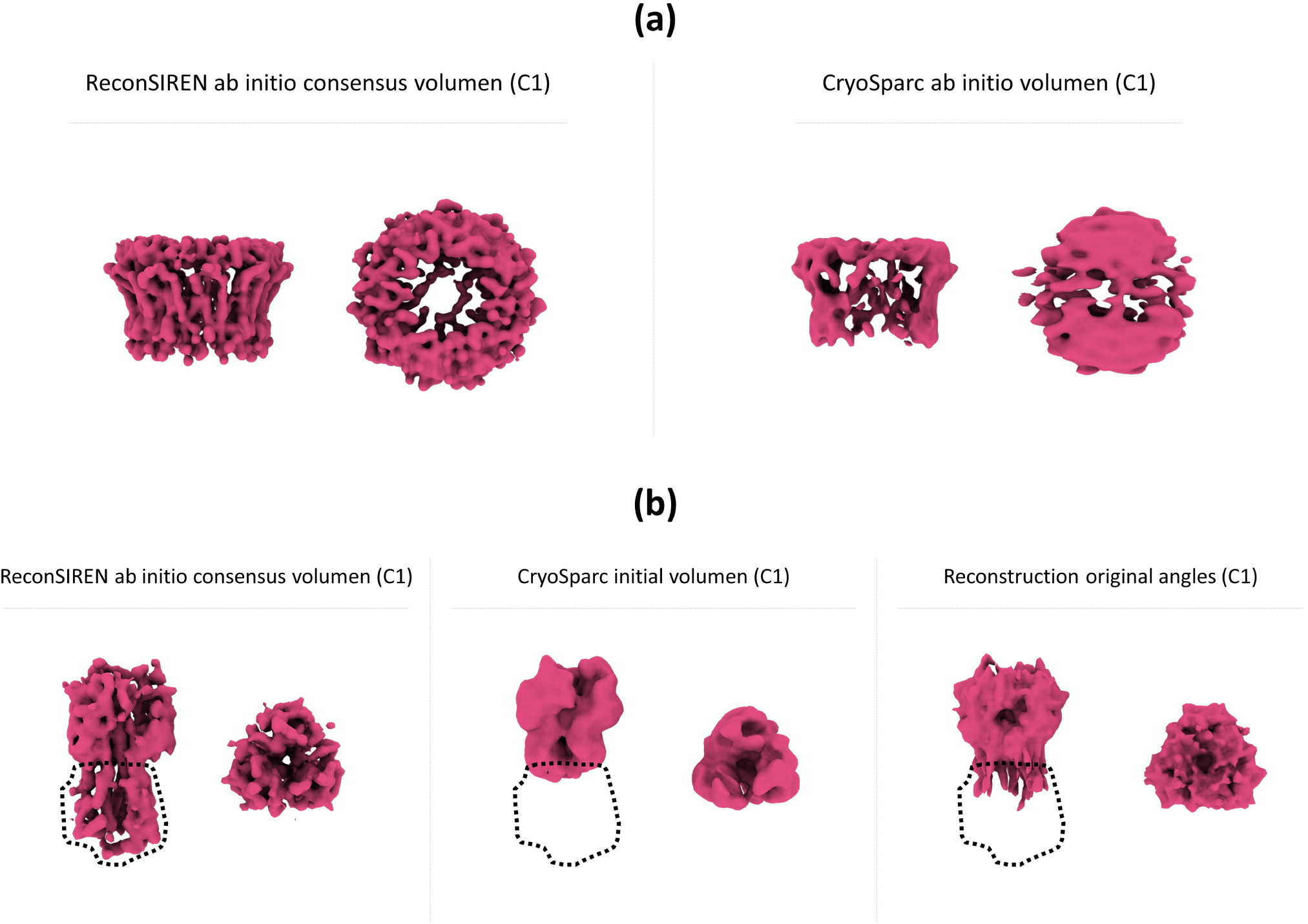
Panel a): Comparison of ReconSIREN and CryoSparc *ab initio* volumes for the CAHRA pose entanglement dataset. Symmetry was not considered in either method. The results show that ReconSIREN produces a high-quality *ab initio* reconstruction even when the protein has high symmetry (C11) and the data exhibit a strong preferred orientation. Panel b): Comparison of ReconSIREN and CryoSparc *ab initio* volumes against the reconstruction of the data published for the hemagglutinin EMPIAR 10096 dataset. Because ReconSIREN avoids attraction artifacts from preferred orientations in the data, the resulting consensus volume recovers the bottom part of the hemagglutinin.

A similar result was observed when analyzing the hemagglutinin dataset as shown in Figure 3b. When comparing the initial volumes from CryoSPARC and ReconSIREN, the CryoSPARC map omits the bottom part of the protein, which is typical for this dataset due to its strong preferred orientation. In contrast, ReconSIREN pose ensemble optimization significantly mitigates attraction, allowing it to recover the protein’s bottom part. Additionally, the Figure compares against reconstructions from the published image alignments and in-plane shifts, showing that the volume also lacks the bottom part of the protein, as expected.

As a final experiment, we further refined the angular assignment estimated by ReconSIREN for the hemagglutinin dataset using a CryoSPARC local refinement job with C3 symmetry. The purpose of this is to check how much a refinement can be improved when the in-plane shifts, angular assignments, and *ab initio* consensus volume estimated by ReconSIREN are used as an initial guess (for local refinement, a smooth mask covering the whole consensus map was used). We show the resulting maps in Figure 4. Because ReconSIREN recovered the bottom part of the protein, we included in the comparison not only the map reconstructed from the untilted dataset, but also the map published with the tilted dataset. The local refinement map starting from the ReconSIREN initial volume recovers the same mass seen in the tilted dataset, with slightly improved structural features. This comparison shows that, even with a strong preferred orientation, it is possible to obtain similar or better structural detail and completeness in the published untilted particles than in the tilted ones when attraction artifacts are absent.

**Figure 4:**
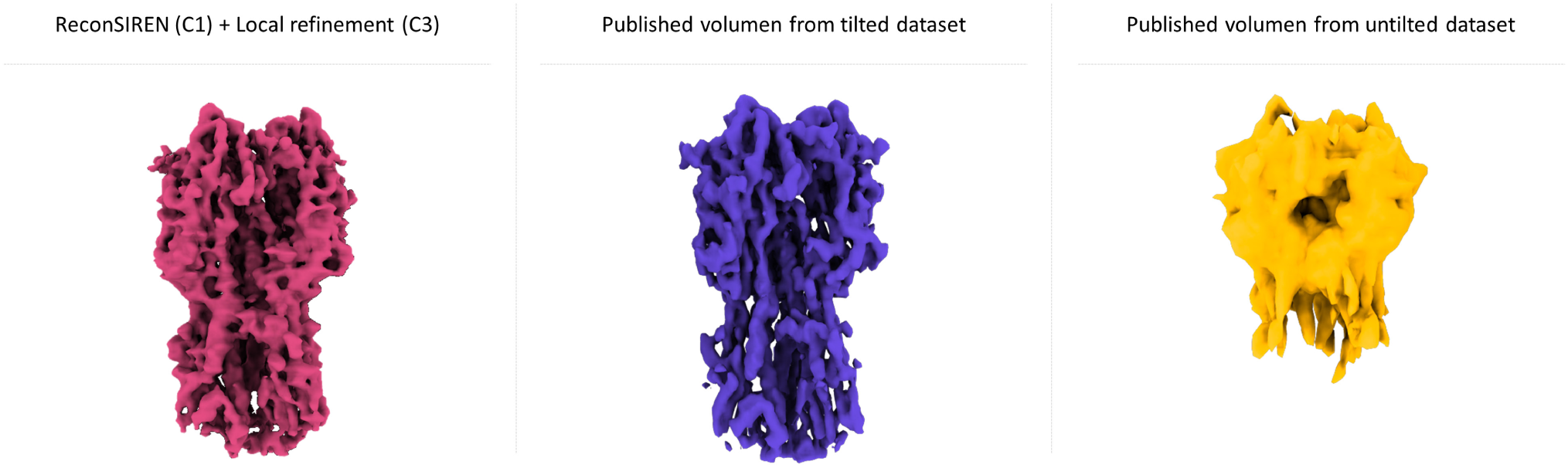
Comparison of the local refined ReconSIREN *ab initio* consensus map against the published volumes for the Hemagglutinin tilted (EMPIAR-10097) and untilted (EMPIAR-10096) datasets. By providing an angular assignment with minimized attraction issues, the refined map obtained from the ReconSIREN results recovers structural features similar to those in the published tilted dataset map.

Overall, these results demonstrate ReconSIREN’s ability to tackle the fundamental CryoEM problem of assigning projection directions correctly without incurring attraction artifacts, even when the data have a strong preferred orientation. This allows ReconSIREN to recover accurate *ab initio* angular assignments and reconstructions even under these challenging conditions.

### Evaluation of ReconSIREN conformational spaces derived from experimental data

Together with the models that predict *ab initio* image alignments, in-plane shifts, and consensus volume, ReconSIREN includes an additional model in its MoE that identifies *ab initio* conformational landscapes and heterogeneous volumes. This allows early analysis of the types of conformational changes in the data (compositional and/or continuous) and extraction of heterogeneous states that can serve as references for later steps in the workflow.

To validate the heterogeneity information from the MoE experts, we present in Figure 5 a sample of the conformational landscapes obtained after training ReconSIREN on the datasets introduced in the previous sections. Note that heterogeneity is not trained sequentially on the image alignments, in-plane shifts, and consensus map; instead, it is trained together with the other modules.

**Figure 5:**
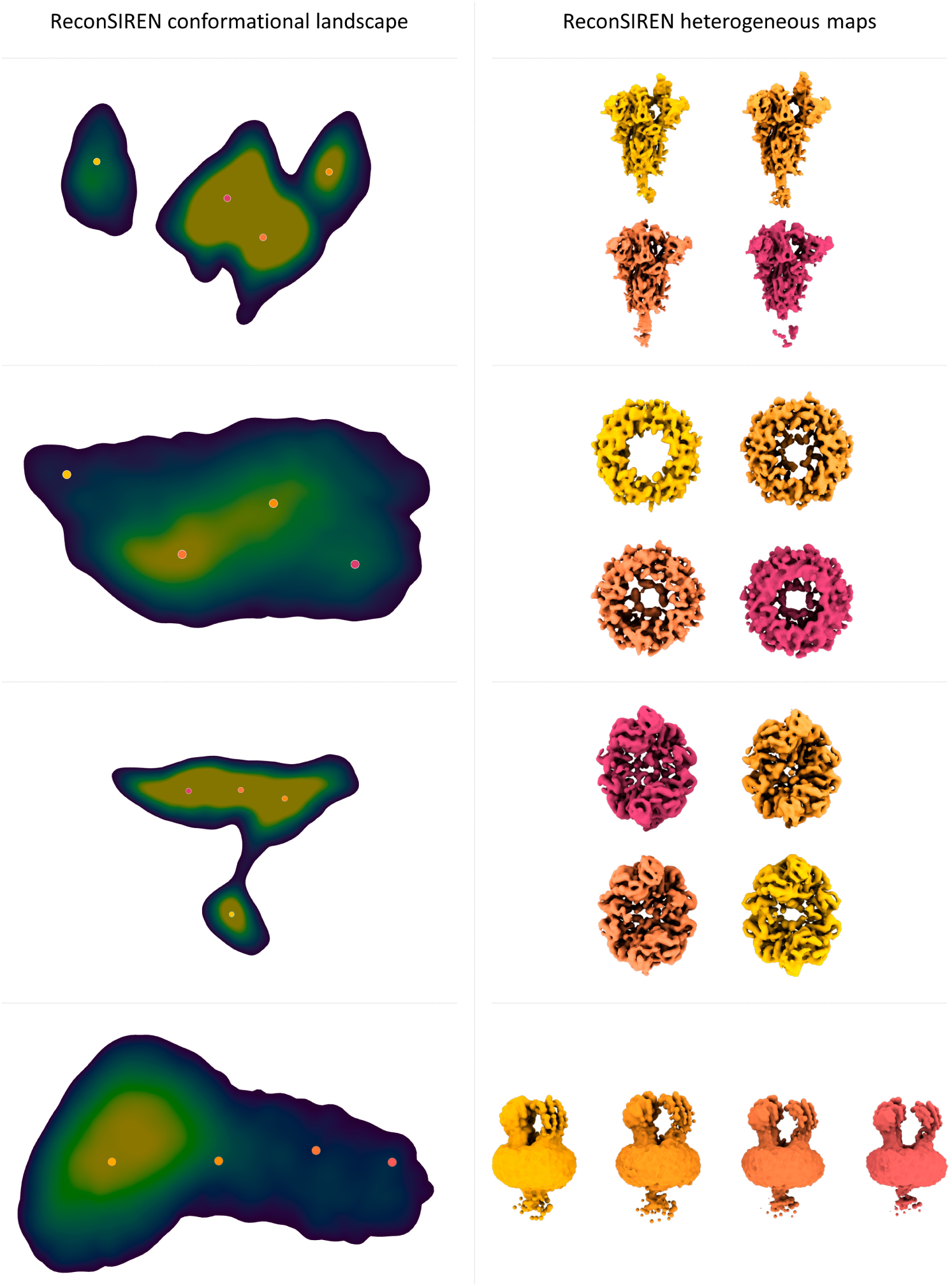
Example of the *ab initio* conformational landscapes and heterogeneous states obtained by ReconSIREN for four different datasets (from top to bottom: SARS-CoV-2 spike, CAHRA pose entanglement dataset, PKM2, ABCB11). The results demonstrate ReconSIREN’s ability to capture compositional and/or continuous heterogeneity early in a processing workflow.

As shown in the Figure, ReconSIREN correctly recovers both compositional heterogeneity (channel dataset) and continuous motions (SARS-CoV-2 RBD spike motion and ABCB11 extracellular domain approximation). We used the PKM2 dataset as a control because it is rigid. The states decoded from the landscape estimated by ReconSIREN for the PKM2 dataset show strong similarity across the reconstructed maps, indicating that the network did not hallucinate motions and correctly detected the protein’s high rigidity, as expected.

Additionally, Figure 6 shows an interesting case with the GR:Hsp90:FKBP51 complex. The protein has two lateral domains that are usually unresolved because of their high intrinsic flexibility. In contrast, ReconSIREN recovers both, as shown in the Figure, allowing us to partially follow their continuous transitions in the conformational landscape. This result, together with the ones presented in Figure 5, validates the ability of ReconSIREN to detect a wide range of structural variation completely *ab initio*.

**Figure 6:**
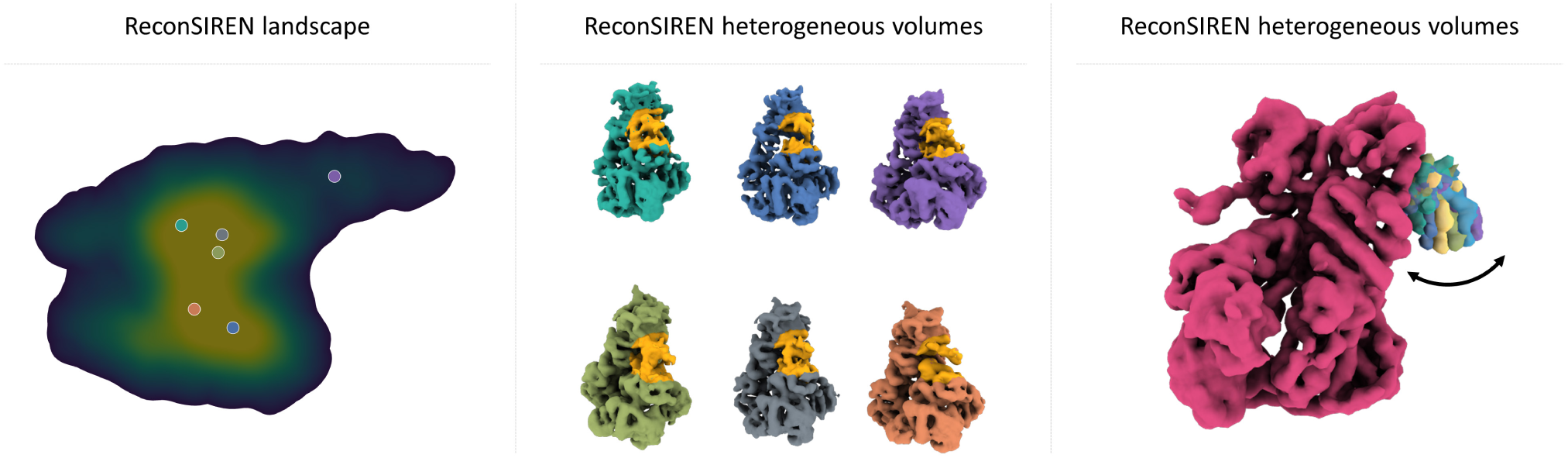
Example of the *ab initio* conformational landscapes and heterogeneous states obtained by ReconSIREN for the GR:Hsp90:FKBP51 dataset. The results primarily show ReconSIREN’s ability to detect highly flexible regions in the GR:Hsp90:FKBP51 complex without prior knowledge of poses or in-plane image shifts.

### Evaluation of ReconSIREN when trained with 2D averages

Another interesting use case of ReconSIREN is its ability to estimate *ab initio* reconstructions from 2D class averages rather than from the initial experimental images. This use case is particularly interesting because the higher SNR of the class averages drives faster convergence than the experimental images. Additionally, ReconSIREN will estimate the angular assignment and in-plane shifts for each 2D average. This information is used, in conjunction with the per-article alignment of each experimental image to its class, to report an initial 3D alignment of each image to the *ab initio* consensus volume. Thus, this per-particle alignment can serve as a starting point for subsequent refinement jobs.

To assess ReconSIREN’s ability to work with 2D averages, we perform 2D classification in CryoSPARC on some of the datasets introduced in the previous sections. We then manually filtered the resulting 2D classes to remove obviously incorrect averages that showed no structure. Figure 7 shows the results after training ReconSIREN with the previous averages.

**Figure 7:**
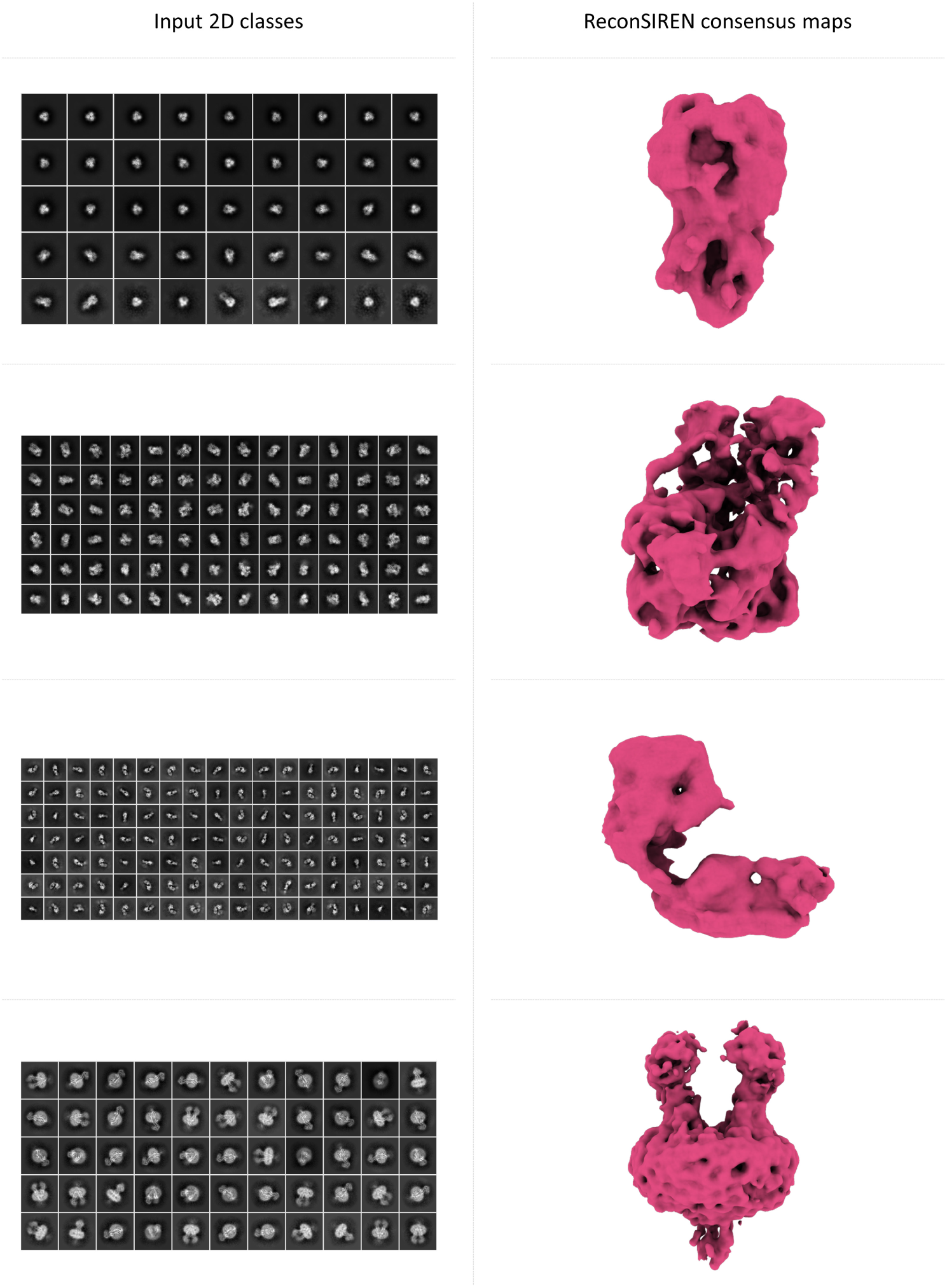
Example of the *ab initio* consensus reconstructions obtained with ReconSIREN for five different datasets (from top to bottom: Hemagglutinin, HSP90, HER2, ABCB11) starting from 2D class averages. By using 2D class averages, ReconSIREN can recover *ab initio* reconstructions with coarse angular assignments with faster convergence and reduced training times.

**Figure 8:**
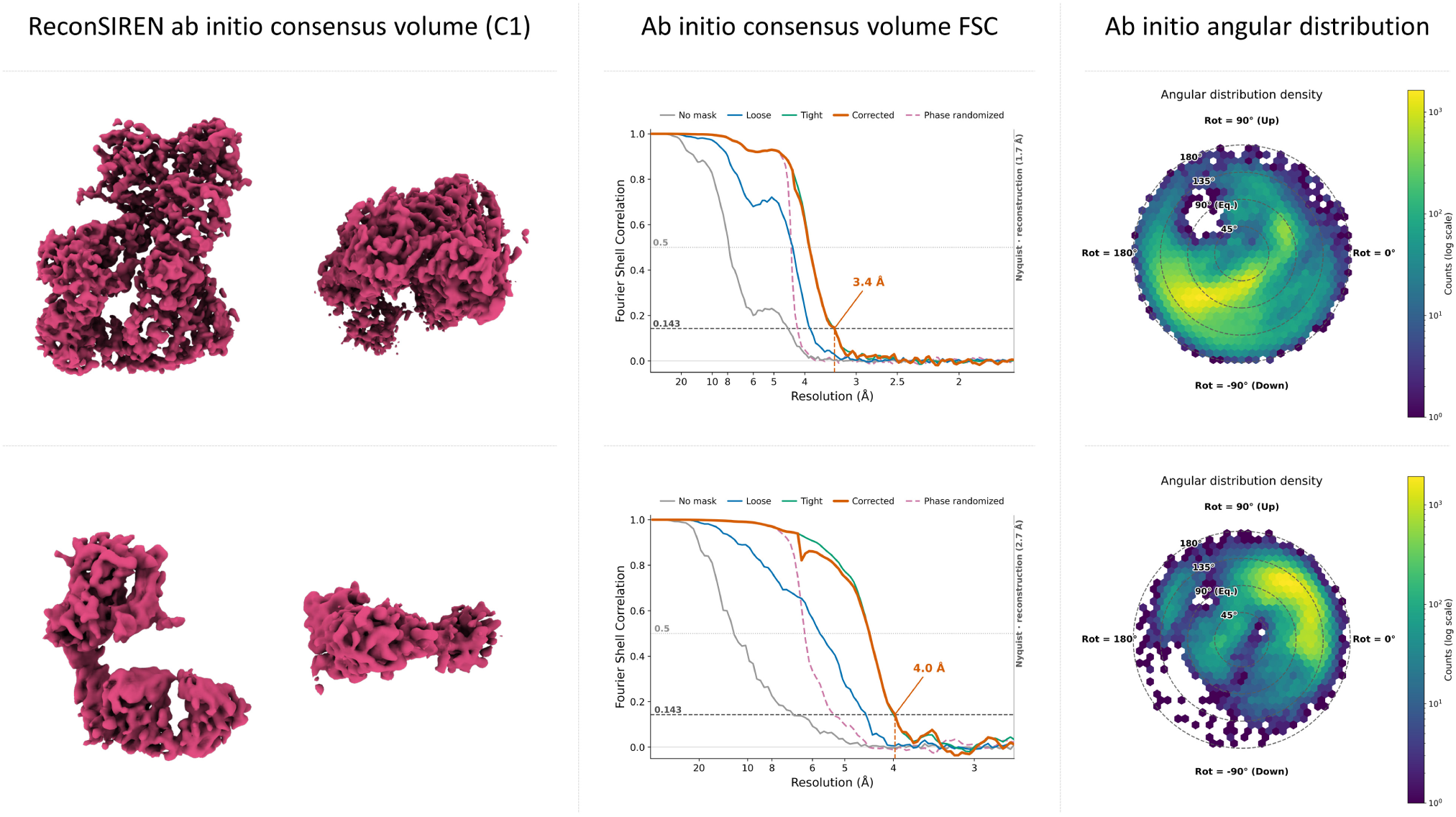
*ab initio* consensus map, FSC curves, and angular distributions obtained for the GR:Hsp90:FKBP51 and HER2 datasets without downsampling with ReconSIREN. The results show that, with full-resolution images, ReconSIREN can produce *ab initio* estimates with quality comparable to, or even surpassing, 3D refinement results, at the expense of longer training times and higher memory usage.

Although the resolution from the averages is not comparable to that from the experimental images, the results show that ReconSIREN can effectively recover high-quality *ab initio* reconstructions directly from the averages. This introduces an additional use case for the method, enabling users to obtain initial volumes more quickly when speed is prioritized over accuracy.

### Analysis of the maximum resolution achievable by ReconSIREN

As shown in previous sections, even when limiting image resolution, ReconSIREN achieves much higher-resolution results than standard *ab initio* methods. This is a property of our network design: ReconSIREN always performs a local search over the parameters it optimizes, so it can refine them on the fly to improve accuracy as they converge.

Therefore, the accuracy achievable in the parameters estimated by ReconSIREN (angular assignments, in-plane shifts, consensus volume, and heterogeneity) is, in principle, not limited to a coarse initial solution, because the network can continue improving them as training progresses by exploring the solution space locally.

Following this idea, we propose a different way to run ReconSIREN, where we directly input full-resolution images to the network. As in the previous experiments, the images carry only their CTF, letting ReconSIREN estimate the remaining parameters fully *ab initio*. Under this setup, we sacrifice some training speed to achieve the highest possible resolution and estimation accuracy.

To illustrate, we ran ReconSIREN on full-size images from the GR:Hsp90:FKBP51 and HER2 datasets. Figure 9 shows the consensus maps, FSC curves obtained with CryoSparc, and the angular assignments obtained after training. The results show that ReconSIREN can achieve *ab initio* estimates comparable to, or even surpassing, results obtained after 3D refinement. Therefore, the resulting volume could serve as sufficient input to the final steps of an image-processing pipeline.

**Figure 9:**
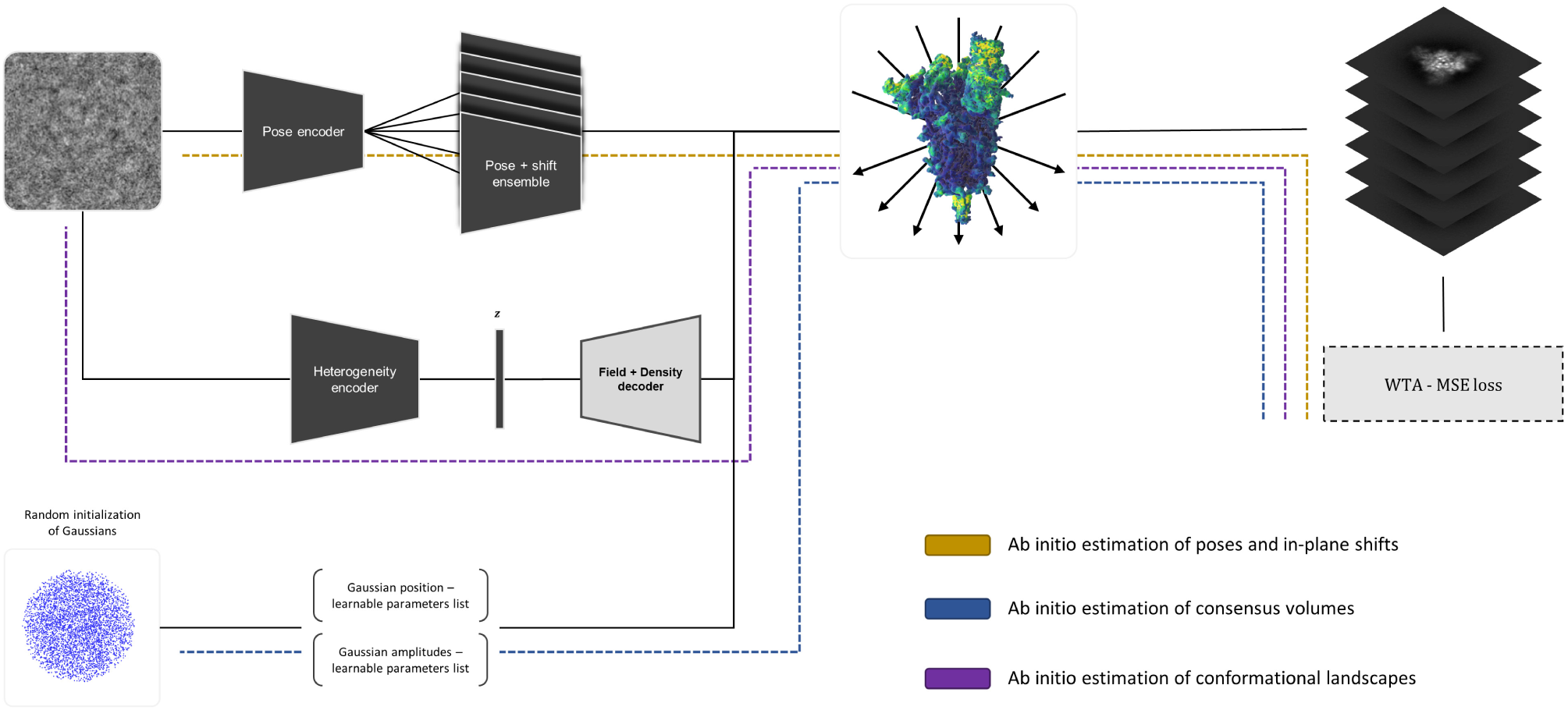
Scheme of the overall architecture of ReconSIREN, including its different independent modules: image alignment module (yellow), consensus volume reconstruction module (blue), and conformational heterogeneity module (purple). ReconSIREN’s modular architecture is self-configurable, allowing activation of the consensus volume reconstruction and conformational variability modules depending on the problem. In addition, it is possible to provide a reference volume **V** to perform a global assignment instead of an *ab initio* reconstruction.

In addition to the previous results, Supplementary Video 1 shows some of the heterogeneous states obtained by ReconSIREN for GR:Hsp90:FKBP51 without downsampling. This dataset is particularly interesting because we already analyzed its heterogeneity with our method HetSIREN [17]. Comparing the two methods, we found that ReconSIREN correctly reports the same kind of heterogeneity as HetSIREN, without needing any prior angular assignment to achieve comparable accuracy.

Overall, the results show that ReconSIREN can achieve high-resolution fully *ab initio* without compromising the accuracy of the different parameters it estimates.

## Discussion

A critical step in any CryoEM image-processing workflow is *ab initio* structure estimation, as well as estimating projection directions and in-plane shifts for the images in a dataset. These estimates provide the initial guesses that later steps in the workflow build on. Therefore, accurate *ab initio* estimates are critical for achieving high-resolution volumes with improved interpretability. This need applies to all specimens studied with SPA methods, but is especially challenging when the specimen shows a so-called preferred orientation, referring to a highly unbalanced projection-space distribution.

The possibility of applying deep learning to these *ab initio* estimates has already been partially explored in methods such as CryoAI [12], CryoSPIN [13], or CryoFIRE [15]. However, most of these methods require *a priori* information to work properly, such as precentering the images to simplify the search for good local minima. They thus are not fully *ab initio*. Additionally, because exploring the projection space with local optimizations is complex, methods in this area have shifted to a hybrid approach that combines classical grid searches with local optimizations to identify the right projections, as CryoDRGN-AI [16] does.

To make deep learning viable for approaching the *ab initio* optimization challenge, we present ReconSIREN, the first neural network capable of tackling the *ab initio* estimation of projection directions, in-plane shifts, and homogeneous/heterogeneous states, even in cases of preferred orientations, relying fully on deep learning principles. We tested ReconSIREN on a wide range of challenging specimens, demonstrating its ability to learn high-resolution *ab initio* reconstructions, as well as highly accurate angular assignments and in-plane shifts. Moreover, the MoE approximation of ReconSIREN enables the network to detect compositional and continuous heterogeneity effectively, even in highly flexible protein regions.

For the initial experiments presented, we downsampled the input images to favor performance over resolution. This explains why some FSC curves report a resolution lower than the Nyquist resolution of the images the network observed. However, because we always computed the FSCs on the original full-size images, the resolution from homogeneous reconstruction can exceed the Nyquist resolution the network observed. If the estimated angular assignments and in-plane shifts are sufficiently accurate, they can further improve resolution when working with non-downsampled images. We also used this experiment to show how to input ReconSIREN results into other software, such as CryoSparc [21], to continue the workflow. ReconSIREN can already output its results as *cs*, *star*, and *xmd* files, making the transition to other packages much simpler.

However, downsampling is not required to train ReconSIREN; it should be used only to speed up training, if desired. Because ReconSIREN always locally optimizes the parameters it searches, it is not, by construction, limited to getting trapped in a coarse solution. Therefore, when full-resolution images are used, ReconSIREN still converges to the right structure and, in some cases, it may achieve results comparable to or even surpassing those obtained after a 3D refinement step. We show this in two examples to highlight the method’s ability to return consensus/heterogeneous volumes that can be used as input to the final steps in an image processing workflow.

In addition, we carefully designed ReconSIREN to directly address a fundamental problem in SPA CryoEM: estimating per-particle angular assignments without attraction artifacts induced by preferred orientations. By testing the network on well-known datasets strongly affected by preferred orientations, we show ReconSIREN’s robustness in learning *ab initio* angular assignments and volume reconstructions while significantly mitigating attraction artifacts. These improved estimates are essential for later steps in the image-processing workflow toward reconstructing high-resolution structural features.

## Methods

The following sections describe the ReconSIREN architecture in detail, focusing on the expert-independent modules that predict angular assignment, shifts, and conformational heterogeneity. After the architectures, we describe the network’s training strategy, including its modular nature and its dependence on the input data.

Figure 9 shows a schematic of the overall network architecture discussed in the next section. To complement this information, Table 1 summarizes the training and prediction times for ReconSIREN, measured on an RTX 6000 Ada Generation GPU.

**Table 1:**
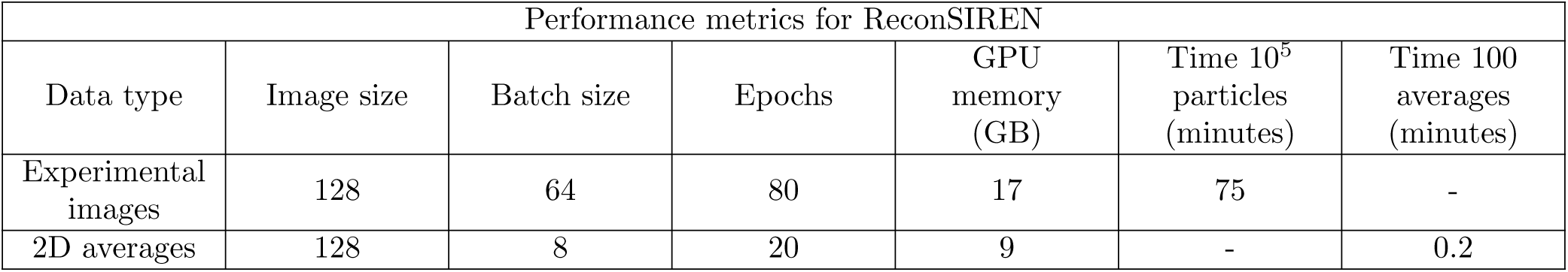
Execution times and GPU memory consumption for ReconSIREN. Metrics refer to the training phase. The reported values correspond to the hyperparameters used to train on all the datasets analyzed and presented in this work.

We acknowledge the partial use of AI tools to test and identify possible bugs during the development of some of the functions in the method presented in the manuscript.

### Architecture of the image alignment module

ReconSIREN is based on an MoE approach, in which different network modules are trained independently using their own cost functions and optimizers to address specific sub-problems that, when combined, can solve a larger problem. We assign each module its own optimizer to customize its training dynamics and improve convergence and training speed.

The first module corresponds to the expert who determines the pose and in-plane shifts of the raw experimental images. This module comprises a common component based on the popular VGG16 [24] architecture, with residual and skip connections among the different VGG16 blocks. In our experiments, we found that skip and residual connections improved both generalization and network training.

One of the main issues of approaching the *ab initio* image alignment problem is the need to explore almost every possible region of the projection sphere.

Many methods address this problem by exhaustively searching a gridded version of the projection sphere, progressing from a coarse to a fine grid to improve the accuracy of the estimated angles. This exploration is essential to ensure convergence; otherwise, the optimization could converge on degraded solutions, typically characterized by strong attractions.

However, deep learning optimization is, by definition, local. Even with advances in deep learning optimizers, optimizing an image’s projection direction may still get trapped in undesirable local minima. To overcome this limitation, ReconSIREN optimizes the projection direction using an ensemble of encoders that predict different possible projection directions for any given image. The ensemble members are initialized so that the projection directions they start optimizing from are evenly distributed on the projection sphere. We call this initialization “the anchor” rotation. Therefore, each ensemble member is, by construction, forced to explore a different region of the projection sphere, reducing exhaustive exploration to N independent local optimization problems. Under this setup, ReconSIREN not only explores the solution space globally but also leverages deep learning’s advanced local optimization to find better local minima.

Another advantage of our approach is its robustness to attraction artifacts, which are usually induced by preferred orientations in the data. Our ensemble approach, combined with a uniform prior over the angular distribution within the ensemble, gives the network a pool of plausible, distinct projection directions to choose from, preventing it from falling into the undesirable solution of assigning the same projection direction to all images due to increased SNR.

At the architecture level, the common encoder output connects to the previous ensemble, consisting of a variable number of MLP-based independent networks (18 by default, unless the user modifies it), which we call heads. Each produces a different pose that, when combined with its anchor rotation, yields the final alignment matrix. To promote an extensive search of the projection sphere, we initialize the kernel of each head to produce small rotations around its anchor.

For in-plane shifts, we use a single MLP head for all images, combined with a delta shift learned by each previous head. We found this approximation converges faster and adds only a minor increment to the memory footprint.

We optimize the previous image alignment model with its own AdamW optimizer and a default learning rate of 1e-4.

Additionally, the image alignment model can refine a precomputed set of image alignments, if available. The user can easily switch between the refinement and *ab initio* modes from the program interface.

### Architecture of the consensus volume reconstruction module

The consensus volume reconstruction model decodes an *ab initio* structure using the poses and in-plane shifts predicted by the previous module.

The module uses a Gaussian Splatting [25] approach to reconstruct the consensus volume from a set of randomly initialized Gaussians within a sphere contained within the volume box. The position and amplitudes of the Gaussians are all learnable parameters organized in the form of a table. This allows independent optimization of the Gaussian displacements and amplitudes needed to reconstruct the consensus volumes. We can express the reconstruction of the consensus volume as follows:

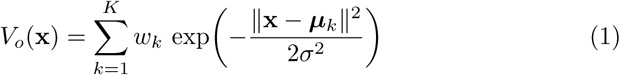

where *K* represents the total number of Gaussians available, *w_k_* and ***µ****_k_* represent the amplitude and center of the k-th Gaussian, and *σ* is the standard deviation. In our case, we set *σ* as a shared learnable scalar for all the Gaussians.

By default, the model initializes 10,000 Gaussians. We found this number to be a good compromise between a high-resolution initial volume and strong performance with a low GPU memory footprint. Additionally, users can also pass an initial consensus volume to the network to initialize the Gaussian positions and amplitudes. When fine-tuning of the initial number of Gaussians is needed, or the default value is too limiting, we recommend starting with two Gaussians per residue and reducing the number if memory becomes a constraint.

The consensus volume model is trained with its own AdamW optimizer at a default learning rate of 3e-3, with global gradient clipping.

### Architecture of the conformational variability module

The last module in ReconSIREN predicts a conformational latent space and the conformational states that originate from the latent-space vectors. This module comprises an independent autoencoder that converts raw experimental images into 3D volumes representing the conformational states captured by the dataset.

The first part of this module comprises a set of convolution layers, followed by a 3-layer MLP that maps raw images into a latent space, thereby organizing them according to their structural information. The encoded latent space vectors are then fed into an SIREN-based MLP decoder with the same architecture as the consensus volume model. The SIREN-based MLP decoder follows the same principles as the one we already introduced in HetSIREN [17], including a new Feature-wise Linear Modulation (FilM) [26]. With this modulation, the network can learn to condition the Gaussian-coordinate input with the latent vectors predicted by the heterogeneity encoder. This new decoder will learn a set of Gaussian amplitudes and displacements such that the volume recovered represents a heterogeneous reconstruction specific to a given image:

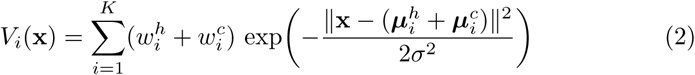

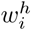 and 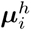 are the Gaussian amplitudes and centers learned by the conformational model from the conformational latent vectors, while 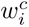 and 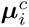 represent the corresponding consensus amplitudes and positions learned by the consensus module.

The conformational model is optimized with its own AdamW optimizer, a default learning rate of 1e-4, and global gradient clipping.

### Handling symmetry in ReconSIREN

Although many datasets can be readily aligned without accounting for protein symmetry, some cases benefit from accounting for a biomolecule’s intrinsic symmetry to improve the quality of both the reconstructed volume and the image poses.

To that end, ReconSIREN internally symmetrizes its predicted poses according to a valid symmetry group. Although this symmetrization affects the image alignment module’s outputs, it also directly affects the volume consensus reconstruction module, forcing the network to learn a reconstruction compatible with the enforced symmetry.

Under symmetry, the angular space ReconSIREN optimizes is not restricted to the symmetric unit, allowing it to obtain a C1 reconstruction directly from the predicted angles. Working in C1 is particularly important for the conformational variability module, as it enables it to handle the asymmetric characteristics of conformational changes.

When symmetry is taken into account, the network randomly generates one valid symmetric projection associated with the predicted poses in the ensemble, forcing it to produce a consensus volume compatible with all symmetric projections as training progresses.

As mentioned earlier, the symmetrization process affects only the consensus volume. Heterogeneous reconstructions obtained from the conformational latent space are forced to have no symmetry, even if the consensus volume has been symmetrized (i.e., the poses seen by the heterogeneous model are never symmetrized). In this way, the delta volumes Δ**V***_i_* predicted by the conformational variability module will learn to recover the correct C1 conformational state from the consensus volume, thereby improving the accuracy of the network’s conformational states.

### ReconSIREN training strategy and cost functions

Thanks to its modular architecture, ReconSIREN can easily configure itself for the problem at hand. The configuration is directly inferred from the input data, determining which modules to activate or deactivate to define the best possible architecture for a given experiment.

To simplify the usage of the program, we restrict the execution of ReconSIREN to the following modes:

- Full *ab initio* mode: Determines from scratch the image angular assignment, in-plane shift, consensus volume, and conformational landscape.
- Refinement mode: Given a set of particles with precomputed alignments and an initial consensus volume, ReconSIREN will jointly optimize both to improve their accuracy. Additionally, ReconSIREN will determine the conformational landscape from the images.
- *Ab initio* angular assignment: Given a fixed consensus volume, ReconSIREN will assign the best projection direction and in-plane shifts to the images, as well as determine their conformational landscape.

To explain how the optimization process of ReconSIREN works, we will focus on the full *ab initio* mode:

1. The forward pass begins by passing the images through the image alignment module, yielding a set of *N* possible image alignments from the head ensembles and the image in-plane shifts.
2. Next, the consensus volume module recovers the consensus volume. This module takes a constant input consisting of the initial Gaussian coordinates and predicts, from these, a refined set of Gaussian centers and amplitudes.
3. The estimated alignments are used in conjunction with the consensus Gaussian positions and amplitudes to render a set of projections from the consensus volume through a Gaussian Splatting approach. The resulting projections will have shapes (*B, N, H, W*), where *B* is the data batch size, *N* is the total number of predicted alignments, and *H* and *W* are the image dimensions. By comparing these images to the experimental ones, we can obtain the representation loss used to train the network:

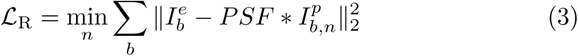 Where 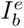 represents a given experimental image in the batch 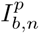, a projection of the consensus volume convoluted with the Point Spread Function (PSF). Based on the previous equation, the representation loss is computed as the hard Winner-Takes-All (WTA) loss over the ensemble dimensions (i.e., only the projection with the lowest Mean Squared Error (MSE) against a given batch image backpropagates through the previous loss). It should be noted that, when a point symmetry group other than C1 is considered, the alignments predicted by the ensemble are further processed by multiplying them by the corresponding symmetry matrices. Instead of considering all possible symmetry matrices for a given point-group symmetry at once, the network chooses one at random and uses it to modify the alignment. This approach recovers a symmetrized consensus volume without additional GPU memory overhead or impact on overall training performance. Also, even when symmetry is taken into account, the network can search over the entire projection sphere. Therefore, the angles predicted by the network do not require a symmetry expansion or a symmetry-breaking step to be used when reconstructing a C1 volume. To discourage the predicted poses from collapsing into a few angular clusters, we penalize the discrepancy between the distribution of viewing directions and the uniform distribution on *S*^2^. Since optimal transport on the sphere is computationally expensive, we estimate this discrepancy using the Sliced Wasserstein Distance. Let **x***_i_* = **R***_i_***e**_3_ ∈ *S*^2^ be the viewing direction of the *i*-th particle, obtained by applying a predicted ensemble rotation **R***_i_* ∈ *SO*(3) to the canonical projection axis **e**_3_ = (0, 0, 1)^⊤^, so that **x***_i_* is simply the third column of **R***_i_* and indicates the direction along which that particle is projected. Given these directions 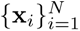, we draw a reference set 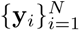 representing a set of uniformly distributed projection directions on *S*^2^ and a set of *L* random projection axes 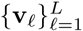 also on *S*^2^. Projecting both sets onto each axis gives the scalars 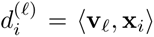 and 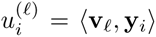 for *i* = 1*, …, N* and *ℓ* = 1*, …, L*, and we compare their sorted values:

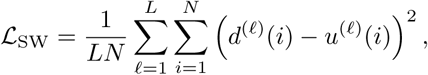

where *d*^(*ℓ*)^(*i*) and *u*^(*ℓ*)^(*i*) denote the *i*-th smallest of the projections along **v***_ℓ_*. The loss *L*_SW_ vanishes precisely when the predicted ensemble directions are uniformly distributed over the sphere. Because **x***_i_* depends on **R***i*, gradients of *L*_SW_ flow back into the pose estimator, which is what allows the term to shape the ensemble angular distribution during training.
4. The last step is to forward the images through the conformational heterogeneity model. By default, the model first maps the images to an 8-dimensional space. The model then decodes the latent vectors to obtain heterogeneous Gaussian positions and amplitudes, thereby recovering a heterogeneous volume. The heterogeneous map learned by the model imposes no symmetry, regardless of the point symmetry used to learn the consensus volume, ensuring that the conformations found are not affected by symmetry. To simplify computing the projections of the heterogeneous volumes, we consider only alignments that pass the WTA criterion from the previous step. Therefore, the resulting projections have shape (*B, H, W*), representing the projections estimated from the best alignments predicted by the pose head ensemble. The reconstruction loss for the heterogeneous model is computed as:

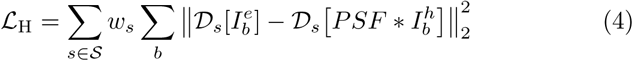 Where 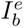 represents a given experimental image in the batch 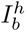, a projection of the heterogeneous volume along the best projection direction determined by the WTA criterion, D*_s_*[·] is a low-pass filter applied to the images (multi-resolution loss), and *w_s_* is a set of weights defining how much each resolution level contributes to the final loss. By default, the loss includes two resolution levels: the original images (weight 0.8) and a downsampled version to 0.6 times the previous box size (but never smaller than 32px) (weight 0.2).
5. The final loss is computed as the sum of the previous losses as:

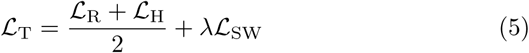

where *λ* is a regularization parameter controlling the strength of the uniform assignment prior.

Each step described above triggers an independent gradient computation based on its losses, optimizing the corresponding module with its corresponding optimizer. We found that this Mixture-of-Experts (MoE) approach converges faster and achieves higher accuracy than training all modules as a single network with a single optimizer.

### Gaussian recycling

One issue we found when tackling *ab initio* reconstruction of consensus and heterogeneous volumes is the loss of expressivity in the Gaussian splatting process when a large fraction of the initial Gaussians are optimized to have near-zero amplitude. These dead Gaussians mainly arise during the first training steps where the network is still exploring the solution space and is far from convergence. This is undesirable because it implies fewer Gaussians will be available to represent the structure once convergence is achieved.

To avoid this problem, we introduced a Gaussian recycling step in ReconSIREN, triggered by default every 5 epochs. During the recycling step, Gaussians having an amplitude below a fraction of the mean amplitude of all the Gaussians in the data are replaced near a randomly selected valid Gaussian under the same criterion. The amplitude of the recycled Gaussian is again set to a fraction of the mean Gaussian amplitude. Thanks to recycling, we ensure that all Gaussians contribute to the optimized structure, maximizing expressivity for a given number of Gaussians.

### Geometry priors for the consensus and heterogeneity modules

To guide the search for consensus and heterogeneous Gaussian positions and amplitudes that represent accurate maps, ReconSIREN includes a series of geometry priors to regularize their optimization.

In order to compute these priors, ReconSIREN estimates the closest neighbor to every consensus Gaussian, as well as the center of mass of the Gaussians weighted by their amplitudes and the radius enclosing 99% of the Gaussians in the consensus. For heterogeneous Gaussians, we also use the closest neighbor, radius, and center estimated from the consensus.

The previous information is used to compute three different geometry priors added as losses during training:

- The first prior penalizes the closest neighbor to a Gaussian whose distance is larger than 2*σ* to promote signal continuity in the map. Additionally, if the closest neighbor is at a distance smaller than 0.7*σ*, it is also penalized, thus preventing the two Gaussians from collapsing in practice to a single one.

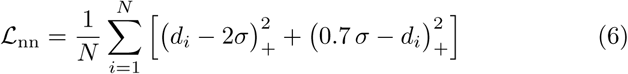 Where *d_i_* represents the distance between a Gaussian and its closest neighbor, and *σ* is the global standard deviation of the Gaussians.
- The second prior promotes that a Gaussian and its closest neighbor have similar amplitude to promote signal continuity at the amplitude level.

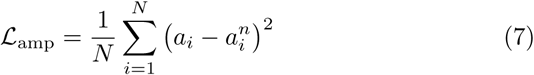 Where *a_i_* represents the amplitude of a Gaussian and *a^n^* the amplitude of the closest neighbor to the previous Gaussian.
- The third regularization penalizes Gaussians that fall outside a sphere whose radius and center are defined based on the previously explained values (center of mass of the Gaussians weighted by their amplitudes and the radius enclosing 99% of the Gaussians in the consensus). This mitigates outliers trying to move away from the map structure.

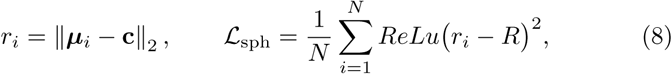 Where *µ_i_* is the mean of a given Gaussian, *c* is the precomputed center of the sphere, and *R* is the precomputed radius of the sphere.

The previous priors are activated by default and usually have a stronger impact when training on full-size images to achieve the highest possible resolution from ReconSIREN.

### Avoiding collapsing of the heterogeneity module

Another issue we found when training ReconSIREN (mainly with full-size images) is that the heterogeneity module can collapse toward the consensus map. This issue arises when the heterogeneity module chases the consensus during training, preventing stabilization and yielding a spurious local minimum where the consensus volume is left as is.

One way to avoid this problem is to start optimizing the heterogeneity module once the other two experts have stabilized. Therefore, ReconSIREN lets users choose which epoch the heterogeneity module should start optimizing to mitigate this issue.

However, it is hard to know a priori when the alignment and consensus modules will stabilize in practice. To make heterogeneity-module optimization robust to collapse, we add two priors at the latent-space level estimated by the heterogeneity module:

- The first prior computes the variance of each dimension in the latent space and promotes that it is not driven towards zero to prevent dimensions from becoming “dead” (i.e., carrying no information at all).

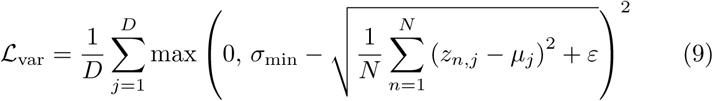 Where *D* represents the total number of dimensions in the latent space, *σ_min_* is the minimum standard deviation allowed for any dimension, *z_n,j_* is the *j* component of the latent space vector of a given image, and *µ_j_* is the mean of one component of the heterogeneity latent space of the network.
- The second prior computes the off-diagonal elements of the covariance matrix of the latent space and drives them towards zero. This encourages the latent-space dimensions to become decorrelated, allowing them to represent independent structural information (i.e., the information in one dimension is not present in another).

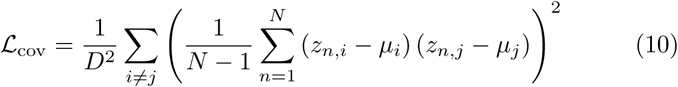 Where *D* represents the total number of dimensions in the latent space, *z_n,i_* and *z_n,j_* are two different components of the latent space vector for a given image, and *µ_i_* and *µ_j_* are the means of two different components of the heterogeneity latent space of the network.

To compute these losses, we use a memory bank to store up to *N* latent vectors from the current batch and previous steps. The memory bank makes it easier to use more samples than in a batch, improving the accuracy of variance and covariance estimates. In the previous equations, the values *µ_i_* and *µ_j_* are computed from the previous memory bank.

## Data Availability

SARS-CoV-2 spike, GR:Hsp90:FKBP51 complex, HER2, and ABCB11 datasets are available upon request. The pose-entangled dataset from the CAHRA challenge is publicly available for download from the challenge’s official webpage. The hemagglutinin dataset is publicly available in EMPIAR (ID 10096). The PKM2 dataset is publicly available in EMPIAR (ID 10648).

## Code Availability

ReconSIREN is freely available as part of the package *hax-em*.

In addition, ReconSIREN can also be used inside Scipion [27] through the plugin *scipion-em-hax*.

## Author Contributions

DH developed and tested the ReconSIREN network presented throughout the manuscript. DH wrote the manuscript. COSS and JMC jointly supervised this work.

## Acknowledgments

The authors acknowledge the financial support from the Ministry of Science, Innovation and Universities (BDNS n. 716450) to the Instruct Image Processing Center (I2PC) as part of the Spanish participation in Instruct-ERIC, the European Strategic Infrastructure Project (ESFRI) in the area of Structural Biology, Grant [PID2022-136594NB-I00] funded by MICIU /AEI/10.13039/501100011033/and “ERDF A way of making Europe,” by the “European Union,” Spanish State Research Agency AEI/10.13039/501100011033, through the “Severo Ochoa” Programme for Centres of Excellence in R&D [CEX2023-001386-S], Comunidad Autónoma de Madrid” through Grant: S2022/BMD-7232, European Union (EU) and Horizon 2020 through grant: HighResCells (ERC - 2018 - SyG, Proposal: 810057) European Union (EU), and Horizon Europe through grant: Fragment Screen Proposal: 101094131. We thank Professor S. Subramaniam and D. Agard for providing us with the spike and GR:Hsp90:FKBP51 datasets processed and presented in this work, as well as for their support and feedback.

---

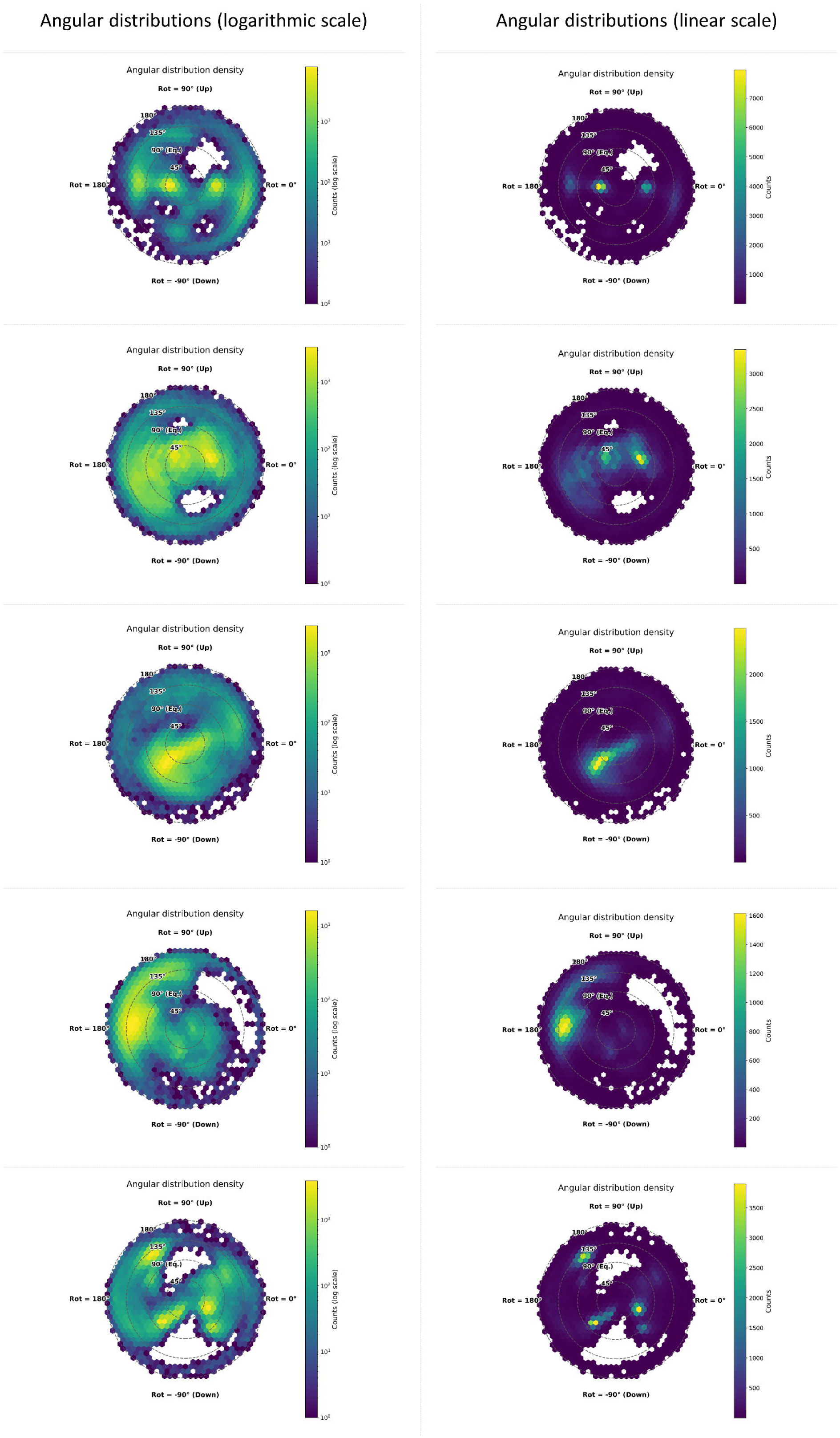

