## Supplementary Information for "*Ab initio* CryoEM maps minimizing preferred orientation problems in heterogeneous mixtures: ReconSIREN"

David Herreros[*1], Carlos Oscar Sanchez Sorzano[+1], and Jose Maria Carazo[+1]

[1]Centro Nacional de Biotecnologia-CSIC, C/ Darwin, 3, 28049, Cantoblanco, Madrid, Spain
[+]These authors jointly supervised this work: C.O.S. Sorzano and J.M. Carazo

* Corresponding author


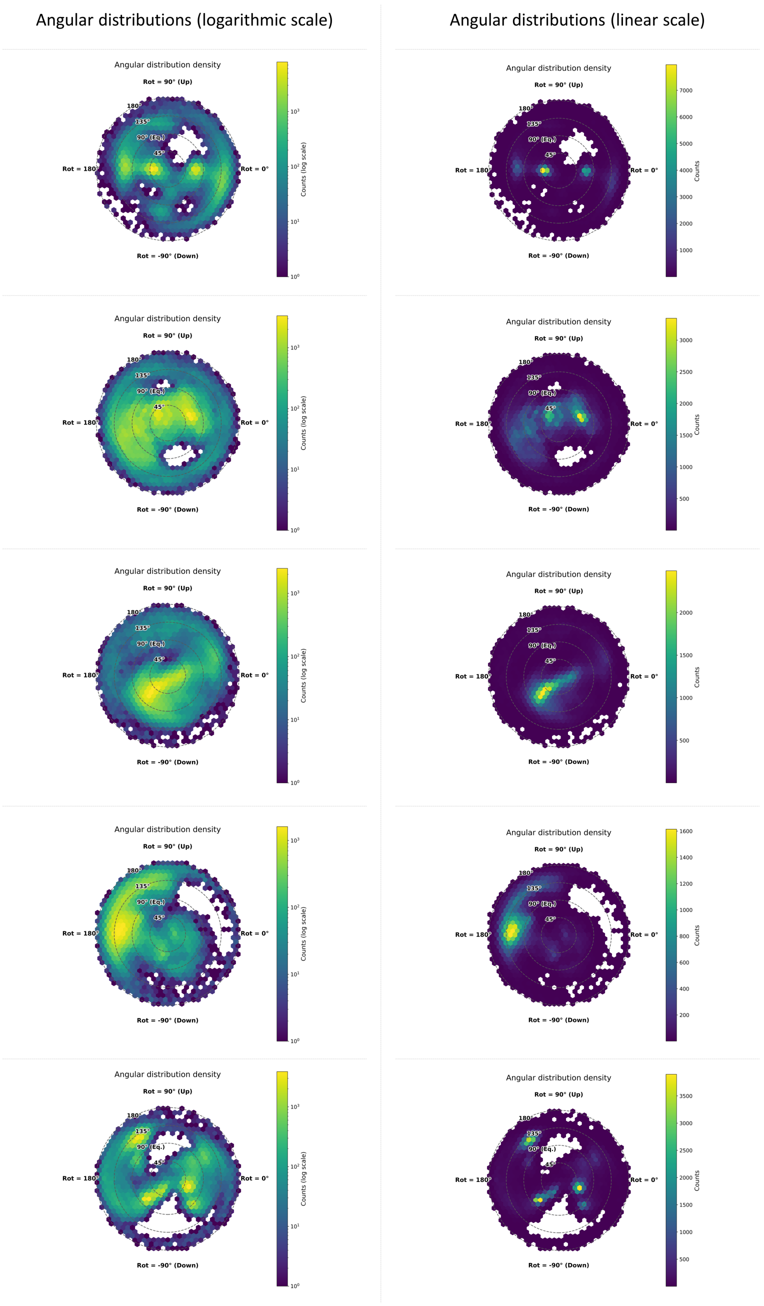


Supplementary Figure 1: Comparison of the angular distributions in logarithmic scale and linear scale. The distributions presented in the paper correspond to the logarithmic-scale distributions, as they visually simplify assessing how the images were assigned in the projection sphere.
